# Interpersonal synchronization of movement intermittency

**DOI:** 10.1101/2021.06.09.447663

**Authors:** Alice Tomassini, Julien Laroche, Marco Emanuele, Giovanni Nazzaro, Nicola Petrone, Luciano Fadiga, Alessandro D’Ausilio

## Abstract

Most animal species group together and coordinate their behavior in quite sophisticated manners for mating, hunting or defense purposes. In humans, coordination at a *macroscopic* level (the pacing of movements) is evident both in daily life (e.g., walking) and skilled (e.g., music and dance) behaviors. By examining the fine structure of movement, we here show that interpersonal coordination is established also at a *microscopic* – *sub*-movement – level. Natural movements appear as marked by recurrent (2-3 Hz) speed breaks, i.e., submovements, that are traditionally considered the result of intermittency in (visuo)motor control. In a series of interpersonal motor coordination tasks, we demonstrate that submovements are not independent between interacting partners but produced in a tight temporal relation that reflects the directionality in the partners’ informational coupling. These findings unveil a potential core mechanism for behavioral coordination that is based on between-persons synchronization of the intrinsic dynamics of action-perception cycles.

## Introduction

Motor behaviour is the product of coordinated patterns of muscle activations which, in humans, can reach a striking level of variety and complexity. To be effective, behaviour needs to be tuned to time-varying changes in the external world. Among the most complex motor activities we are capable of, many arise from a fairly spontaneous tendency to synchronize our movements to periodic or quasi-periodic stimuli (1) or to other people’s movements (2-5), providing the basis for perhaps the richest forms of social interactions that exquisitely belong to our species – i.e., playing music and dancing (6).

Interpersonal coordination can indeed be incredibly smooth and accurate, in particular if it is rhythmically organized. Such a capacity has been the object of intense investigation over the last two decades, either by looking at spontaneously emergent patterns of coordination during natural behaviours (7) (e.g. walking side by side, hand clapping, postural sway, limb swinging) or, typically, by asking participants to tap jointly according to simple beats or more complex musical rhythms (8, 9). Many metrics have been put forth to quantify synchrony as well as tempo (co-)adaptation in individual (1, 10) and joint (4, 11) rhythmic performance, showing that, even without any specific training, people are capable of pacing their movements with near-millisecond accuracy.

However, by zooming into the fine structure of movement, another form of rhythmicity becomes apparent at a lower level. Movement is in fact organized into smaller units – *submovements* – which combine together to make up full kinematic trajectories (12, 13). If observed with the appropriate lens, movement is never really smooth but marked by kinematic discontinuities that are clearly appreciable at slow speeds. Such discontinuities, or speed breaks, are not a consequence of biomechanical constraints, nor an incidental or erratic phenomenon but tend to recur with specific periodicity in the range of 2-3 Hz. The presence of submovements has first been noticed more than a century ago (14); since then submovements have been documented in many studies, in human (13, 15, 16) as well as non-human primates (17-20), and generally ascribed to an intermittent control of movement, deemed as an efficient computational strategy to face inherently long sensorimotor delays (12, 21-24).

Each submovement is thought to consist in a pre-programmed (open loop) motor correction that can only be issued in an intermittent fashion. To some extent, their presence (13) and properties [e.g., periodicity (20, 25, 26)] depend on the available visual feedback of movement. In particular, artificially increasing feedback delays during visuomotor control tasks (such as hand tracking of a visual target) alter submovement frequency in a consistent manner (20). Yet, submovements seem not to be just a mere consequence of extrinsic delays in the visuomotor loop. Recent evidence highlights how the intrinsic oscillatory dynamics within the motor system contributes as well (19, 27, 28), and in a way that is partly independent from (experimentally manipulated) feedback delays (20). Submovements may thus arise as a consequence of oscillatory neural dynamics reflecting visuomotor control loops (20).

Interpersonal coordination requires continuous corrections based on an accurate (and mostly visually mediated) estimation of others’ behaviour towards a joint motor outcome. For such coordination to be successful, information must be flowing within both individual and inter-individual action-perception loops. Movement intermittency opens an empirical window upon these action-perception loops, hence providing the opportunity to explore the fabric of how behavioural visuomotor coordination is mechanistically established.

Here, we exploit a movement synchronization task that has been classically used to probe interpersonal coordination (4) but we purposely focus on the faster time scale of movement intermittency. We show that interpersonal synchronization does occur also at a lower level than that pertaining to the sequencing and pacing of movements, i.e., the level of submovements. Importantly, the timing of submovements clearly captures a functional coupling between partners, showing that movement intermittency is actively co-regulated during the interaction.

## Results

Sixty participants forming 30 couples performed a movement synchronization task. As shown in Figure 1A,B, participants were asked to keep their right index fingers pointing towards each other (without touching) and perform rhythmic flexion-extension movements around the metacarpophalangeal joint as synchronously as possible either in-phase (towards the same direction) or anti-phase (towards opposite directions). We instructed participants to keep a slow movement pace (i.e., with each flexion-extension cycle lasting about 4 s) by having them practicing in a preliminary phase with a reference metronome set at 0.25 Hz (metronome was silenced during task performance). Each participant also performed the same finger movements alone (solo condition) with the only requirement of complying with the instructed pace (i.e., 0.25 Hz). Finger movements were recorded using retro-reflective markers tracked by a 3D real-time motion capture system (Vicon), providing with continuous kinematic data sampled at 300 Hz (Figure 1C).

**Figure 1.**
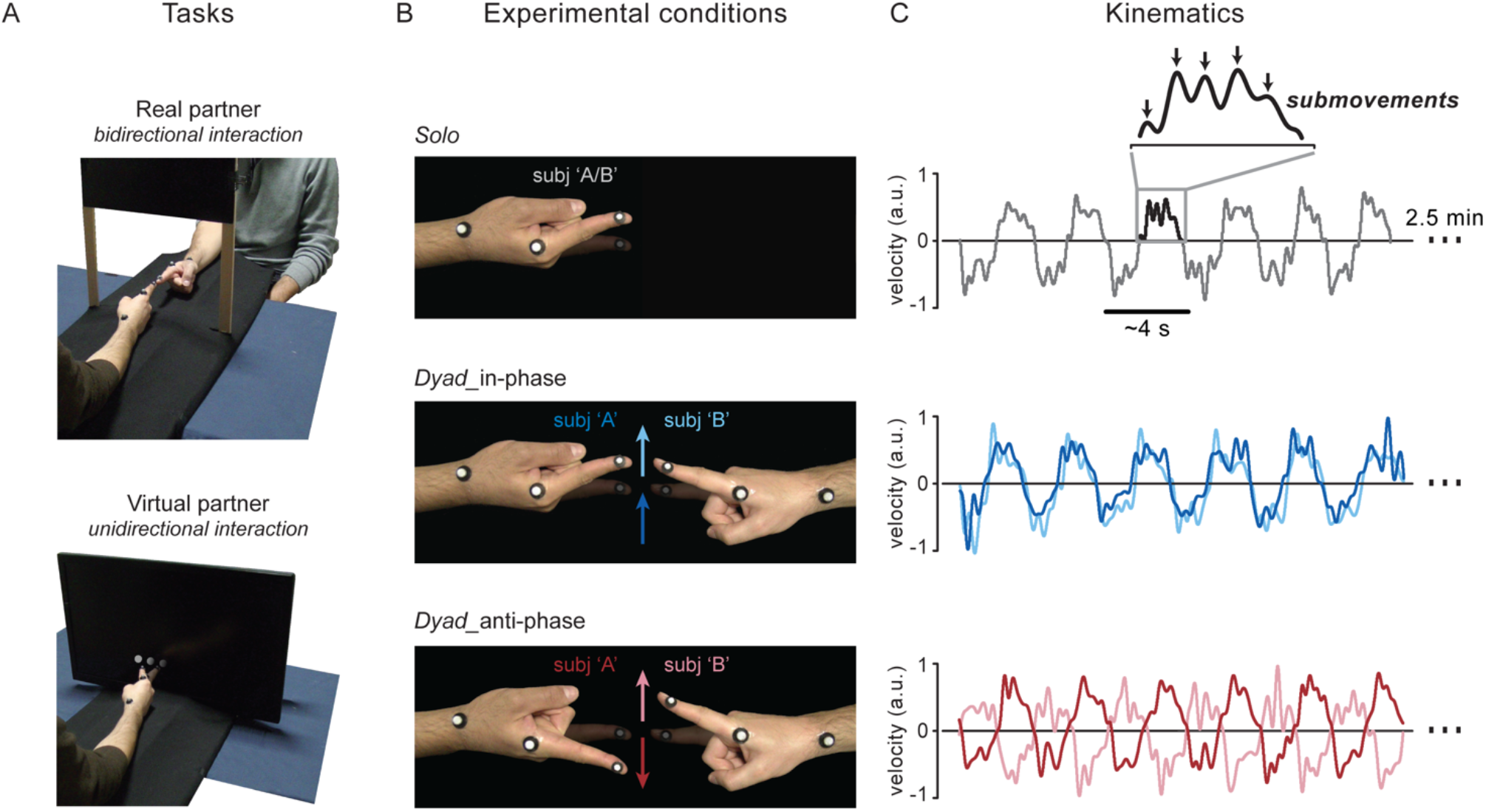
Experimental setup and procedure. **(A) Tasks**. Main task (‘Real partner’) and one of the three secondary tasks (‘Virtual partner’; for the other tasks see Supplementary Figure 2). In both tasks, participants seated at a table with the ulnar side of the right forearm resting on a rigid support and performed rhythmic (0.25 Hz) flexion-extension movements of the index finger about the metacarpophalangeal joint. In the main task, participants formed couples (n = 30) and were asked to synchronize their movements to one another (dyadic condition; top panel); whereas in the ‘Virtual partner’ task, participants (n = 20) synchronized their movements to a visual dot that moved on a screen according to a pre-recorded human kinematics (bottom panel). **(B) Conditions**. Finger movements were performed by each participant alone (solo condition; top panel) as well as together with the (real/virtual) partner (dyadic condition; the middle and bottom panels illustrate the ‘Real partner’ task). In the dyadic condition, participants were required to keep their fingers pointing straight ahead without touching each other (or the screen) and move as synchronously as possible either in-phase (towards the same direction; middle panel) or anti-phase (towards opposite directions; bottom panel). **(C) Kinematics**. Movements were recorded in blocks of 2.5 min (2 blocks per condition) using a real-time 3D motion capture system (Vicon; sampling rate: 300 Hz). Examples of the participants’ finger velocity along the main movement (x-)axis, measured at the distal phalanx of the index finger (see markers in Figure 1A), are shown for all conditions. Periodic (2-3 Hz) submovements are highlighted in the inset.

### Rhythmicity at movement and submovement levels

By task design, movements are rhythmically organized with periodicity very close to the instructed pace. To highlight the rhythmic components of movement, we transformed the velocity time series into the frequency domain. As expected, all conditions display a major spectral peak around 0.25 Hz – referred to as F0 (Figure 2A). Moving together with a partner leads to a general speed-up (29, 30) as shown by the F0 frequencies being higher than the instructed movement pace for both dyadic conditions (in-phase: 0.28 ± 0.037 Hz, p <0.001; anti-phase: 0.27 ± 0.037 Hz, p = 0.002), whereas slightly lower for solo performance (0.24 ± 0.036 Hz, p = 0.059; mean ± SD; one-sample t-tests against 0.25Hz; see Figure 2A). No difference in pace is, however, observed between in-phase and anti-phase synchronization (p = 0.445; paired samples t-test).

**Figure 2.**
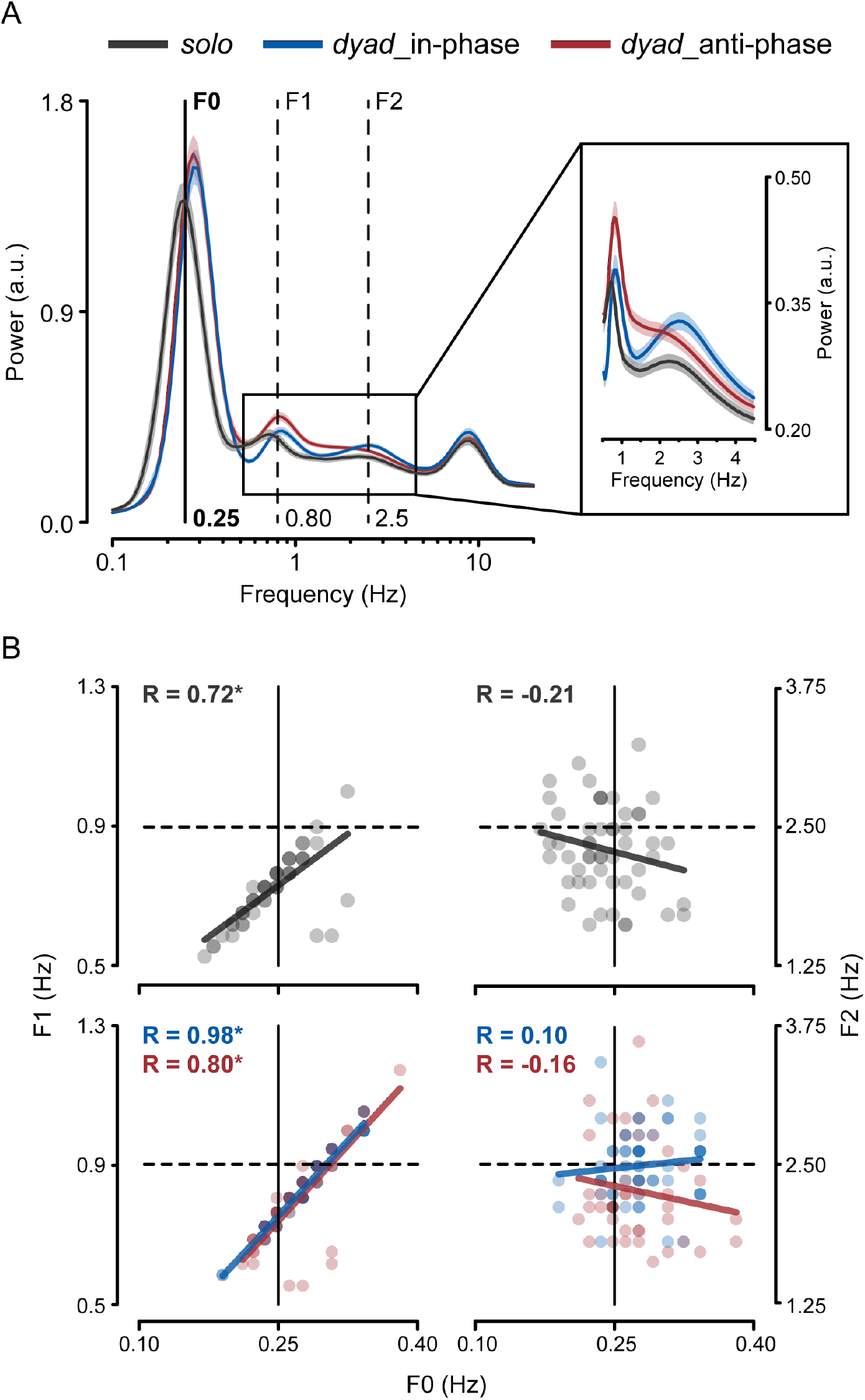
Rhythmicity at movement and submovement levels. **(A)** Power spectrum of finger velocity for all conditions (solo, dyad-in-phase/anti-phase; mean ± SEM). The main spectral component peaking around the instructed movement rate (i.e., 0.25 Hz) is denoted as F0 (black solid line). The spectral components peaking around 2.5 Hz (submovement-related) and around 0.8 Hz are denoted as F2 and F1, respectively (dashed lines), and highlighted also in the inset. **(B)** Scatter plots showing (across-subjects) correlations of F0 peak frequencies with F1 (left) and F2 (right) peak frequencies for the solo (top) and dyadic (bottom) conditions. Data points represent individual participants.

Besides the obvious F0 rhythmicity, the velocity profiles are marked by regular pulses that recur every ∼300-500ms (see Figure 1C for an example), yielding a less prominent but distinct spectral peak also in the 2-3 Hz range – indicated as F2 (Figure 2A). These pulses – otherwise called submovements – are a basic kinematic feature and are indeed present irrespective of the specific coordination mode. However, the increase in power at 2-3 Hz appears to be more sharply defined (narrow-band) for in-phase than anti-phase synchronization (see inset in Figure 2A), suggesting that submovements may be produced more regularly when subjects are engaged in the former than the latter type of coordination.

Importantly, the F0 and F2 peak frequencies are not correlated (across-subjects), neither for solo performance (R = -0.21, p = 0.12) nor for in-phase (R = 0.10, p = 0.43) and anti-phase (R = -0.16, p = 0.29) coordination, indicating that submovements periodicity does not hold harmonic or other (linear) relationships with the actual pace of the movements (Figure 2B, right column; note that subjects lacking clear peaks in the velocity power spectrum were excluded from the correlation analyses, see Methods).

The velocity power spectrum shows two additional components. One – denoted as F1 – peaks at about 0.8 Hz for dyadic coordination and slightly lower (∼0.7 Hz) for solo performance (Figure 2A). In sharp contrast with the submovements-related component, F1 is strongly and positively correlated with F0 in all conditions (solo: R = 0.72, p <0.0001; in-phase: R = 0.98, p <0.0001; anti-phase: R = 0.80, p <0.0001; Figure 2B, left column). Because of its tight association with the rate of movement, F1 is of little interest in relation to movement intermittency. Finally, finger velocity shows a faster component with similar spectral properties across conditions and center frequency of ∼8 Hz, which is known as physiological tremor (31).

### Partners synchronize at both movement and submovement level

To quantify coordination at multiple timescales during dyadic interaction, we computed the frequency-resolved phase-locking value (PLV, i.e., a measure of consistency in the phase relationship) between the two partners’ finger velocities. Remarkably, interpersonal synchronization does not occur solely at F0 (and related F1) frequency as prompted by task instructions, but also at F2, the frequency of submovements (Figure 3A). That is, the two partners’ submovements happen to be in a stable (phase) relationship. Yet, this is not true for all conditions. Indeed, submovement-level synchronization is present during in-phase coordination – as shown by the distinct F2-peak in the PLV spectrum – but nearly absent during anti-phase coordination.

**Figure 3.**
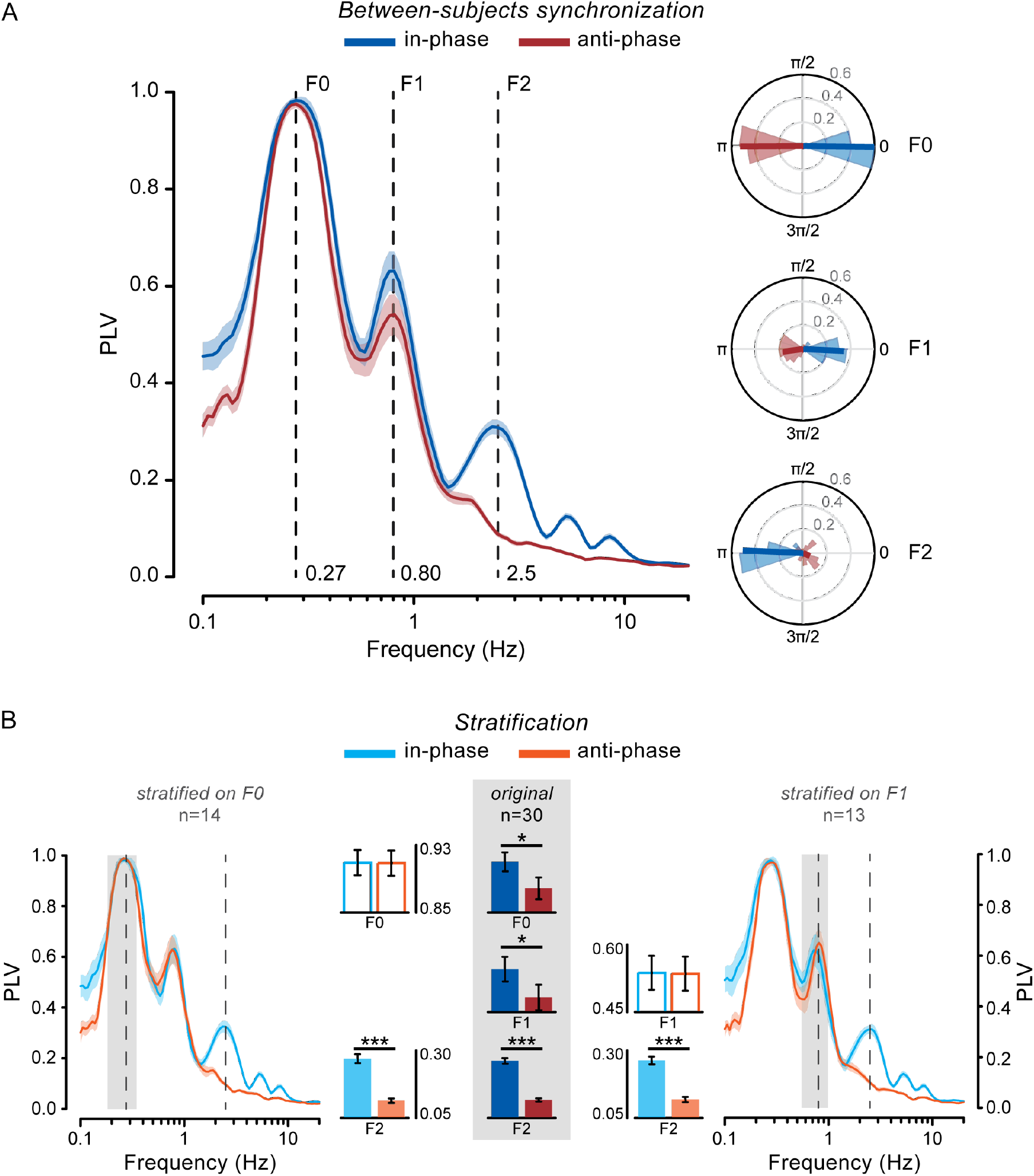
Partners synchronize at both movement and submovement level. **(A)** Between-subjects PLV spectrum for the in-phase and anti-phase condition (left; mean ± SEM). Polar plots (right) showing the across-couples distribution of the mean phase difference (phase lag) for F0 (top), F1 (middle) and F2 (bottom). **(B)** PLV spectra after data stratification (at the group level) on F0 (left) and F1 (right). Bar plots show mean PLV in the relevant frequency ranges for the original (middle; n = 30) and stratified (left, n = 14; right, n = 13) data. Error bars indicate ± SEM. *p<0.05, ***p<0.001.

We ascertained that the observed synchronization at 2-3 Hz is not artefactual or just trivial – e.g., a mere consequence of doing the same movements at the same time – by performing different control analyses. In principle, the 2-3 Hz discontinuities could be produced in a similar way on each movement, thus retaining a consistent phase across movements. Such a phase-locking of submovements to movement onset is actually very modest and comparable for the in-phase and anti-phase condition (see Supplementary Figure 1). Yet, because participants are moving simultaneously, submovements could appear as if they were synchronized between the two partners only by virtue of their (albeit weak) phase-locking to each partner’s individual movement. The greater synchrony between the two partners’ movements during in-phase compared to anti-phase coordination, evidenced by the correspondingly higher PLV at both F0 (p = 0.028) and F1 (p = 0.019; see bar plots in Figure 3B), would further explain why (fictitious) submovement-level synchronization emerges only in the former and not the latter condition. To ensure that submovement-level synchronization is not a by-product of the overall better in-phase than anti-phase (movement-level) synchronization, we used two complementary analytical strategies. Both of them are aimed at matching the conditions (in-/anti-phase) for movement-level synchronization, but the first analysis does so at the group level whereas the second one at the couple level. We first used a data stratification approach that consists in subsampling the couples (by means of a random iterative procedure) so that the mean PLV at F0 and (in separate runs) at F1, is equated as much as possible between the two conditions (see Methods). As shown in Figure 3B, both types of data stratifications leave the pattern of results virtually unchanged, with synchronization at submovement frequency (F2) being significantly stronger during in-phase than anti-phase coordination (p<0.001; independent samples t-tests). As for the second analysis, we computed again the between-subjects PLV, this time not on the entire velocity time series (shown in Figure 3A) but on shorter 2-s segments (covering approximately the duration of one single movement) that are time-aligned to the onset of each partner’s individual movement. In this way, we artificially compensated for the temporal asynchronies between the two partners’ movements, levelling the discrepancy in synchronization performance between the two conditions. Remarkably, this alignment-on-movement procedure severely disrupts synchronization at 2-3 Hz for the in-phase condition, making it as weak as that observed for the anti-phase condition (p = 0.215; paired-samples t-test; see Supplementary Figure 1). Altogether, these results decisively exclude that the observed phenomenon is explained by any systematic locking of submovements to movement dynamics and/or the better in-phase vs. anti-phase movement synchronization performance; rather, they suggest a true and real-time coadaptation of submovements between the two partners.

Interpersonal synchronization at submovement frequency is confirmed also in two additional experimental conditions. In the first one, we changed hand posture and movement axis (from horizontal to vertical) and demonstrate that the effect is independent from the congruency between the two partners’ flexion/extension movements during in-phase and anti-phase coordination (see Supplementary Figure 2A). In the second one, we show that a similar phenomenon persists when the task involves whole-arm movements (on the horizontal and vertical plane), thereby extending our observations to multi-joint action coordination (see Supplementary Figure 2B).

Thus, submovement-level synchronization seems to be largely independent from the effector as well as the congruency between the partners’ movements, but highly dependent on their (visuo-)spatial alignment. In fact, the main difference between the two coordination modes is the spatial alignment of the effectors’ endpoints (i.e., the fingertips): a 0- and 180-deg difference in position is required for in-phase and anti-phase coordination, respectively. Consistently with task requirements, the (across-couples) distribution of the mean phase lag at F0 between the two partners’ kinematics is strongly concentrated around 0-deg for in-phase and 180-deg for anti-phase coordination (Figure 3A, polar plot on top). The same phase relationship as for F0 is also observed for the related F1, although with less degree of consistency. Surprisingly, synchronization at submovement-level (Figure 3A, polar plot on bottom) is characterized by an opposite, ∼180-deg, relative phase shift with respect to what is established at movement-level (F0) during in-phase coordination (phase lags are relatively scattered during anti-phase coordination, in agreement with the PLV lacking a distinct F2 peak in this condition). In other words, submovements during in-phase synchronization are interlocked in the two partners and seem to follow one another with an alternating, counterphase, pattern.

### Bidirectional and unidirectional (co-)modulation of submovements

To further clarify the nature of submovement-level synchronization, we computed the cross-correlation between the two partners’ (unfiltered) velocities. We first selected data segments that correspond approximately to single movements, i.e., from movement onset to mean movement duration (see Methods). To discard the main contribution deriving from slow and movement-locked components, we subtracted from each segment the mean velocity profile over all segments. We then computed the (between-subjects) cross-correlation either by keeping both partners’ data aligned to the individual movement onset (as just described) and thus misaligned in time (movement-alignment), or by (re)aligning one of the two partners’ data to the other partner’s movement onset (subject ‘A’ by convention) and thus preserving their real alignment (time-alignment). The cross-correlation profile shows a striking difference between the two types of alignments, which is far most apparent for the in-phase condition (Figure 4A). Correlation for the movement-aligned data is maximal at lag zero and slowly declines for lags up to ±0.6 s, reflecting residual (not accounted for by the average subtraction) covariation in movement dynamics between the two partners. Conversely, correlation for the time-aligned data is relatively low at lag zero but sharply increases at symmetrical lags of about ±0.18 s. This curious double-peaked correlation profile most certainly reflects the succession in the partners’ submovements. Indeed, the two cross-correlation peaks are separated by a (lag) interval of almost 0.4 s, closely matching the oscillatory period of submovements production (i.e., 2.5 Hz). While the two peaks are evident in the in-phase condition, two flattened humps are barely detectable at longer lags of about ±0.25 s in the anti-phase condition (Figure 4A, right), reflecting the fact that generation (Figure 2A) and interpersonal locking (Figure 3A) of submovements occurs rather erratically and at a slightly lower frequency in this coordination mode. The rhythmic 2.5-Hz co-variation between the two partners’ kinematics during in-phase coordination and its impairment during anti-phase coordination is further emphasized by taking the difference between the two cross-correlations (time-vs movement-aligned) which yields, for the first condition, an oscillating profile, whereas for the second, an almost flat profile (see Figure 4B).

**Figure 4.**
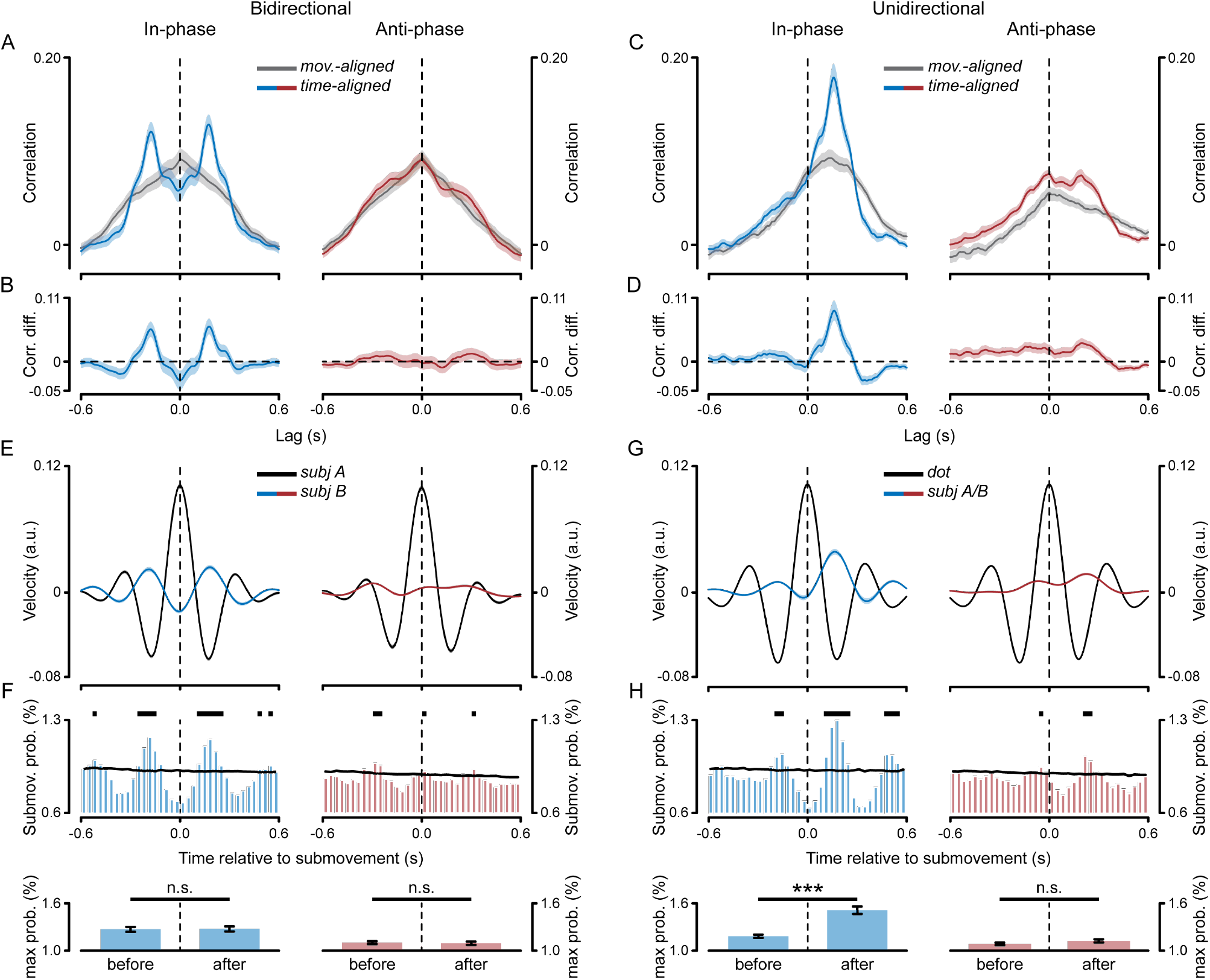
Bidirectional and unidirectional (co-)modulation of submovements. **(A)** Cross-correlation between the two partners’ (unfiltered) velocities during in-phase (left) and anti-phase (right) synchronization in the ‘Real partner’ task (bidirectional interaction). The cross-correlation is computed between velocity data segments (∼2 s) that are either movement-aligned, i.e., aligned to each partner’s movement onset, or time-aligned, i.e., aligned to one of the two partners (subject ‘A’ by convention) movement onset, thus preserving their real time alignment (mean ± SEM; see Methods). **(B)** Difference between the time- and movement-aligned cross-correlation profiles (mean ± SEM). **(C)** Cross-correlation as shown in (A) but computed between the participants’ velocity and the dot velocity in the ‘Virtual partner’ task (unidirectional interaction; note that the dot is used as the reference signal for the time-alignment; mean ± SEM). Correlation at positive/negative lags indicate that the participants’ (sub)movement follows/precedes the dot (sub)movement. **(D)** Same as in (B) but for the ‘Virtual partner’ task. **(E)** Velocity for both partners – subjects ‘A’ and ‘B’ – locked to submovements generated by one partner in the couple – i.e., subject ‘A’ by convention (‘Real partner’ task; mean ± SEM). **(F)** Submovement probability for one participant (subject ‘B’) as a function of time from submovements generated by his/her partner (subject ‘A’; top panels). The black lines indicate the 95% confidence intervals based on surrogate distributions (see Methods); the black bars indicate the time points that survive permutation statistics (Bonferroni-corrected for multiple comparisons across time). Maximal submovement probabilities (for subject ‘B’) computed separately before and after time zero (i.e., subject ‘A’ submovements; bottom panels). **(G)** Same as shown in (E) but obtained by locking the velocity to the dot submovements (‘Virtual partner’ task; mean ± SEM). **(H)** Same as shown in (F) but obtained by computing the participants submovement probabilities as a function of time from the dot submovements (time zero). Error bars indicate ± SEM. ***p<0.001.

If the two partners’ submovements do alternate with regularity (in-phase condition), by locking the velocity of one participant to his/her own partner’s submovements, the probability of observing a submovement in the former should not be uniform over time but significantly higher than chance level at specific times (corresponding to the lags of maximal cross-correlation and to half period of the submovements frequency). Figure 4E,F shows that this is exactly what we observe. We identified submovements as velocity peaks occurring within the movements performed by only one of the two participants in the couple (again subject ‘A’ by convention) and then segmented both participants’ velocities based on the identified peaks (from -0.6 to +0.6 s; see Methods). The submovement-locked velocity for subject ‘A’ shows the expected peak at time zero and two smaller peaks at about ±0.35 s, reflecting 2-3 Hz periodicity in submovements generation. Most interestingly, subject ‘B’ velocity (locked to subject ‘A’ submovements) also shows an oscillating pattern, which is apparent for the in-phase and less so for the anti-phase condition (Figure 4E). The probability of observing a submovement (i.e., a velocity peak) in subject ‘B’ is clearly modulated as a function of time relative to his/her partner’s submovements generation (time zero), and significantly exceeds that obtained for shuffled data (chance level) at multiple and regularly interspersed time points, closely matching the submovements rate (Figure 4F; see Methods). Analogously to what reported for the cross-correlation profiles, subject ‘B’ submovements probability is maximal at relatively shorter (±0.18 s) and longer (±0.3 s) times for in-phase and anti-phase, respectively, suggesting faster (besides tighter) between-subjects alternation of submovements in the former than the latter condition.

The alternating pattern of submovements is likely the result of an inherently bidirectional interaction whereby a motor correction (submovement) in one participant ‘triggers’ another one in his/her partner and so on. But are submovements really consequential to one’s own partner submovements? To put this hypothesis to test, we made the interaction to be no longer bidirectional but purely unidirectional. Ten couples (among the original sample) were also asked to synchronize their movements (in-phase and anti-phase) to a virtual ‘partner’ – a visual dot shown on a computer screen – which was yet moving according to a pre-recorded human kinematics of the same finger flexion/extension movements (see Methods and Figure1A). Remarkably, the timing of the participants’ submovements is tightly related to that of the dot submovements. However, both the cross-correlation (Figure 4C,D) as well as the submovement-locked (Figure 4G,H) profiles are marked by a highly asymmetrical pattern which contrasts with the symmetrical one observed for the real interaction (Figure 4A,B,E,F). Specifically, for the in-phase condition, submovement probability is systematically higher at times following than preceding the virtual partner’s submovements (p <0.0001, Figure 4H) whereas it is symmetrical before and after the real partner’s submovements (p = 0.854, Figure 4F); though with a clear reduction in the strength of submovements modulation, a similar trend can also be observed for the anti-phase condition (virtual partner: p = 0.061, Figure 4H; real partner: p = 0.635, Figure 4F; paired samples t-tests on maximal probabilities before vs. after time zero; see Methods). The reported pattern exactly fulfils what is expected based on the bidirectional and unidirectional nature of the motor corrections in the real and virtual interaction, respectively. To ascertain that submovements are truly co-modulated in a bidirectional way during the real online interaction, we examined the results also at the single couple level. In fact, the observed (group-level) symmetry might conceal asymmetrical results in individual couples (e.g., subject A’s submovements influencing to a larger extent subject B’ submovements than vice versa) that are mixed up in the average due to the arbitrary assignment of subject A/B. Yet, significant asymmetry in the cross-correlation and in the submovement-locked profiles is only found in 3 couples out of 30 (paired samples t-tests on maximal correlation/velocity before vs. after lag/time zero; see Methods), indicating that a bidirectional co-regulation of submovements is indeed occurring in the great majority of the couples.

## Discussion

Whereas an extensive literature looks at the ‘macroscopic’ structure of interpersonal rhythmic coordination – i.e., the sequencing and pacing of movements – the present study looks into its ‘microscopic’ structure – i.e., the intermittency in movement. We show that interpersonal synchronization is not only established at the instructed movement pace – as commonly described (4, 8) – but also at a faster timescale and lower level of the motor control hierarchy, the one reflected in the production of submovements.

Submovements are a general, long-known feature of movement (12). They are particularly evident during sustained and visually guided movements – hand tracking is indeed the task of choice for studying them (13) – but their presence is almost ubiquitous in motor behaviour as shown in a huge variety of tasks and effectors (32-35). Early (12, 21, 22) and recent theoretical accounts (23, 24) mostly consider submovements to be the behavioural sign of underlying intermittency in the control of movement [but see also (36) for an account based on dynamic motor primitives]. Central to this motor control framework is the idea that measured (or estimated) visual errors are used to update the motor commands in a temporally sparse and not continuous manner, that is, only at discrete moments in time. Once updated, motor commands are executed in a feed-forward fashion, eventually building up (seemingly smooth) movement from a concatenation of motor corrections, so-called submovements. The frequency of submovements production (and possibly that of motor updating) has been variably explained based on either intrinsic, time-dependent, factors [e.g., refractory periods, (37, 38) and neural motor dynamics, (19, 20)], or extrinsic and task-dependent factors, such as delays (20, 25, 26), amount (18, 39) and reliability (23) of actual, estimated and/or prediction errors, and most often a combination of both factors (20, 24). However, irrespective of the ongoing debate on the theoretical and computational account of submovements, it is widely agreed that submovements represent a behavioural proxy for the closing of (individual-level) visuo-motor control loops.

When interpersonal movement coordination is demanded, visuomotor control cannot however be closed-up within a single individual. The control loop must be extended to incorporate feedback related to one’s own as well as one’s partner action. In other terms, the relevant visual error signal becomes a joint product of self’s and other’s movements. Crucially, in this case – and not when tracking an external moving stimulus – similar visuomotor machineries are simultaneously at work in the two interacting agents and may thus act in concert. Previous research focusing on joint motor improvisation has considered individual kinematic discontinuities as jittery motion marking epochs of poor coordination performance (2, 40). However, the relational rather than individual control policy of submovement generation was completely overlooked. Strikingly, the present results show that the two partners’ submovements are actually interdependent and time-locked to one another.

Several pieces of evidence indicate that the reported phenomenon does reflect an active co-regulation of submovements between the interacting partners. Submovement-level coordination depends upon the movement-level coordination mode, being it manifest during in-phase coordination but largely impaired during anti-phase coordination. This difference persists regardless of hand posture, and thus of whether the two partners activate simultaneously homologous muscles (flexors/extensors); nor it is a trivial consequence of the greater between-partners synchrony that is observed in this as well as in prior work [e.g.,(30, 41)] during in-phase vs anti-phase coordination. It rather descends most likely from the visuospatial constraints proper to the two modes of coordination. Indeed, moving anti-phase requires a (position-dependent) visuospatial transformation that is instead unnecessary when moving in-phase. This presumably translates into noisier, more unreliable and perhaps delayed computation of visual errors, all known to be important factors in shaping movement intermittency (18, 23, 26, 39). Notably, Reed & Miall (2003) (42) have shown that progressively increasing the spatial separation between the displayed target and the hand cursor correspondingly reduces intermittency (and accuracy) of tracking, probably by making visual evaluation of positional errors coarser and thus less effective in driving feedback-based motor corrections. Analogously, if interpersonal submovements synchronization is mediated by common access to a mutually controlled visual error signal, we would expect it to be degraded during anti-phase coordination because of the inherent difficulty posed by the 180-deg alignment in measuring that very same error. In engineering jargon, anti-phase coordination would be deemed to entail a larger ‘error dead zone’, a concept borrowed by motor control theories to denote the range of inputs to which a system is unresponsive, i.e., which does not bring an update of the (motor) control signals (39). Thus, results differences between in-phase and anti-phase coordination are compatible with functional constrains stemming directly from feedback-based motor control mechanisms.

Yet, a major aspect of our results is that submovements alternate in the two partners, i.e., they are produced in a seemingly counterphase relationship. This clearly excludes spurious or trivial forms of ‘passive’ (zero-phase) coupling and rather points to an ‘informational’ coupling whereby submovements are consequential and reciprocal adjustments to the partner’s behaviour. Indeed, submovement probability is highly non-uniform and maximal ∼200ms before as well as after one’s partner submovements. This temporal symmetry denounces a bidirectional influence which is consistent with the fact that we did not assign explicitly nor manipulate implicitly the partners’ roles (e.g., leader/follower). Results on individual couples are largely consistent with the average effects, showing that implicit leader-follower roles were – if ever – spontaneously assumed by very few couples (i.e., 3 out of 30). Instead of manipulating the partners’ roles, we here replaced one of the two partners with a virtual, unresponsive, partner (a dot moving according to a pre-recorded human kinematics). In this case – i.e., when the interaction is made by design to be purely unidirectional – the symmetry is clearly broken. Now, the participants’ submovements more often follow rather than precede the dot submovements, indicating they are effectively ‘triggered’ by the discontinuities in the dot motion but – obviously – not vice versa. These results decisively corroborate a functional account of interpersonal submovement coordination, making a strong case for a true ‘causal’ relationship between the two partners’ (ongoing) submovements. Thus, although appearing as subtle features, kinematic discontinuities are nevertheless systematically read out and, most importantly, used for coordinating with other people’s movement.

We can argue that coupling of movement intermittency in the two partners originates from corresponding coupling of the underlying visuo-motor loop dynamics, though this hypothesis needs further support from physiological data. Many studies examining coherence between brain activities and kinematics have focused on higher-level, explicit movement dynamics (e.g., movement pace) during solo (43) and interpersonal coordination (44). Remarkably, recent human (20, 27, 28) and monkey (19, 20, 45, 46) evidence has also pointed to specific neural markers of submovement generation. The spectral fingerprint of such neural activity matches well with the periodicity of submovements [i.e., delta/theta-band: 2-5 Hz (19, 20, 28)]; further, this motor activity is phase-locked to submovements (19, 20, 27, 28) and shows consistent dynamics that is unaltered by artificial changes in feedback delays (20). Intrinsic rhythmicity of neural motor dynamics may thus map directly onto the intermittency of motor control.

Computational and physiological perspectives on movement intermittency have usually attributed it solely to the motor stage: intermittent planning and execution of motor commands is based on otherwise continuous feedback reading (and prediction). Movement-related dynamical changes in sensitivity – e.g., sensory attenuation/suppression – are however very well documented phenomena (47). Not only, an increasing bulk of evidence shows as well that sampling of sensory data may routinely operate in a discontinuous, rhythmic fashion (48). Such rhythmicity is often captured by corresponding fluctuations in neuronal excitability and perceptual performance, and it is interpreted as being of attentional origin (49-51). However, most recent work suggests that rhythmic sensory sampling can also be specifically coupled to motor behaviour/activity [(52-57); for reviews see (58, 59)]. Notably, fluctuations in visual perception are not only synchronized to eye movements (60-62) but also to rhythmic cortical dynamics subtending upper limb motor planning (54) and continuous control (63). Although continuous processing has been called in to explain the fast spinal and transcortical reflexive responses (64), intermittency could indeed represent a fundamental property of the more complex sensory and motor functions underlying voluntary behaviour, providing an efficient way for synchronizing time-consuming processing within the action-perception loop.

In conclusion, we show that movement intermittency is effectively synchronized between interacting partners. Such synchronization is likely to constitute an important building block of the low-level visuomotor machinery underlying interpersonal movement coordination. The present investigation opens up a new window upon the (neuro)behavioural mechanisms enabling joint action coordination. This mechanism can be expected to be of crucial importance whenever the interaction poses an accuracy requirement based on the computation of (visual) errors between one’s own and others’ actions (e.g., passing a small/fragile/dangerous object) as well as when learning new motor skills by imitation of others’ behaviour.

Collective or joint action coordination is perhaps the hallmark of human sociality, and a great deal of research is currently pursuing the exploration of its developmental trajectory, comparative origin as well as its relation to pathological conditions (e.g., psychiatric, neurological). The present work has taken a different approach and revealed some core mechanisms of motor control and how they come into play in producing complex coordinated behaviour.

## Methods

### Subjects

Sixty participants (30 females; age 23.9 ± 4 years, mean ± SD) formed 30 gender-matched couples. Except one author (A.T.), participants were all naïve with respect to the aims of the study and were paid (€12.5) for their participation. All participants were right-handed (by self-report) and had normal or corrected-to-normal vision. The study and experimental procedures were approved by the local ethics committee (Comitato Etico di Area Vasta Emilia Centro, approval number: EM255-2020_UniFe/170592_EM Estensione). Participants provided written, informed consent after explanation of the task and experimental procedures, in accordance with the guidelines of the local ethics committee and the Declaration of Helsinki.

No power analysis was used to decide on the sample size (i.e., the number of couples). Sample size estimation was based both on pilot studies as well as on previous studies investigating rhythmic interpersonal synchronization (30, 44) and submovements in visuo-manual tracking (20).

### Setup and procedure

All couples (n=30) performed the same main task (‘Real partner’) and also one of three different secondary tasks (randomly assigned), so that each secondary task was completed by 10 couples in total. Task details are described in the following.

#### Main task – Real partner

Participants were seated at a table either alone or in front of each other (∼1 m apart) with their faces hidden from view by means of an interposed panel (Figure 1A). They were asked to keep the ulnar side of the right forearm resting on a rigid support and perform rhythmic flexion-extension movements of the index finger about the metacarpophalangeal joint (Figure 1A, B). Movements were performed by each participant alone (solo condition) and by the two participants together (dyadic condition). In the latter condition, participants were required to keep their index fingers pointing straight towards each other (without touching) and move as synchronously as possible either in-phase (towards the same direction) or anti-phase (towards opposite directions, Figure 1B). Given the mirror-like participants’ arrangement and hand posture, in-phase coordination required them to perform simultaneously different movements (i.e., as one participant performed finger flexion, the other performed extension and vice versa). Conversely, anti-phase coordination required the two participants to perform the same type of movement (either flexion or extension; see Figure 1B).

In all conditions (solo, dyad-in-phase, dyad-anti-phase), participants were instructed to keep their movement rate around 0.25 Hz (15 bpm; flexion-extension cycle: 4 s). Before the experiment, they familiarized themselves with the reference rate by listening to a metronome and synchronizing their movements to the auditory beat for a short time (∼1 min). Metronome was also played prior to each experimental condition for a few seconds; during the recording blocks the metronome was silenced, and movements were thus self-paced.

#### Secondary tasks

Ten couples completed a second task (Real partner – Hand prone) that was similar to the main one, except that they kept their right hand in a prone posture and moved the index finger along the vertical rather than horizontal axis (Supplementary Figure 2A). As opposed to the main task, during in-phase coordination the two participants had now to perform simultaneously the same type of movement (either flexion or extension), while during anti-phase coordination they had to perform different movements (as one performed flexion, the other performed extension and vice versa).

Other 10 couples completed a secondary task (Real partner – Arm) involving movement of the whole forearm, primarily around the elbow joint (Supplementary Figure 2B). Arm movements were performed (block-wise) along either the horizontal or vertical inner dimensions of the window (∼40×40 cm) delimited by the interposed panel. Only two conditions were tested: participants moved alone (solo) or tracked each other’s movement by keeping the respective fingertips spatially aligned (dyad-in-phase; dyad-anti-phase was not tested in this task). Given the larger movement amplitude, the instructed movement rate was reduced to 10 bpm (i.e., ∼0.17 Hz).

The remaining 10 couples completed a secondary task (Virtual partner) whereby hand posture and finger movements were exactly the same as in the main task. However, instead of coordinating together, participants were now asked to track a visual dot (size: 1.5 cm, position: 7 cm above the bottom screen edge) moving horizontally on a computer screen in front of them (see Figure 1A). The dot velocity corresponded to the velocity of the index fingertip recorded on author A.T. while she was taking part to the main task. The displayed kinematics belonged to all the three tested conditions (2 blocks per condition, see below) – i.e., solo, dyad-in-phase and dyad-anti-phase. Participants tracked the dot kinematics taken from solo performance both in-phase as well as anti-phase; in contrast, the dot kinematics taken from the in-phase and anti-phase conditions were only tracked by the participants in the corresponding modes, that is, in-phase and anti-phase, respectively. Results are shown only for the dot kinematics taken from solo (Figure 4C, D, G, H) as this provides an unbiased comparison between conditions (same displayed kinematics but tracked in different modes); anyhow, results obtained for the other displayed kinematics are qualitatively comparable and support the same general conclusions.

### Kinematic data recording

Movements were recorded along three axes (mediolateral, X; anteroposterior, Y; and vertical, Z) using a ten cameras motion capture system (Vicon; sampling rate: 300 Hz). Three retro-reflective markers were placed at the following anatomical locations on the right hand: on the distal phalanx of the index finger (marker diameter: 6.4 mm), on the metacarpophalangeal joint (marker diameter: 9.5 mm) and on the styloid process of the radius (marker diameter: 9.5 mm; see Figure 1B).

The kinematics recorded on author A.T. and used in the ‘Virtual partner’ task corresponds to the x-axis velocity component of the marker attached on the fingertip (distal phalanx). A photodiode (1 × 1 cm) was placed in the bottom right corner of the monitor and was used for accurately aligning the participants’ recorded kinematics with the displayed dot. A white square was drawn on the screen at the position of the photodiode (hidden from view) in synchrony with the start of the dot motion. The signal from the photodiode was acquired with the same system used to record the kinematic data (i.e., Vicon; sampling rate: 1800 Hz).

### Data collection

Each couple completed the whole experiment in ∼1.5 h. For all tasks, data were collected in separate recording blocks with short pauses in-between blocks. Two blocks of 2.5 min each were recorded for each condition (solo, dyad-in-phase/anti-phase) in all tasks except the ‘Arm’ control task (vertical and horizontal) for which three blocks of 1.5 min were recorded for each condition (solo, dyad-in-phase). The two/three blocks per condition were always performed in succession. Instructions about task/condition were provided verbally before each block sequence. Tasks and conditions order were randomized across couples.

### Data analysis

Analyses were performed with custom-made Matlab code and the FieldTrip toolbox (http://www.fieldtriptoolbox.org; RRID:SCR_004849).

Analysed data corresponds to position data along the main movement axis for the marker attached on the fingertip. Velocity has been computed as the first derivative of position and normalized (block-wise) on maximal speed.

#### Spectral analysis

Spectral analysis was performed by band-pass filtering (FIR filter, order: 3 cycles, two-pass) the continuous (2.5 min) velocity time series applying a sliding window along the frequency axis (range: 0.1-20 Hz) in 100 steps and bandwidths (range: 0.01-3 Hz) that were logarithmically (log10) spaced.

Frequency-resolved instantaneous power was derived by means of the Hilbert transform and then averaged over time points and blocks (Figure 2).

The between-subjects phase-locking value (PLV) was computed as the mean resultant vector of the instantaneous phase differences (Hilbert-derived) between the two partners’ velocity time series (the resulting PLV was then averaged across blocks; Figure 3).

The (between-subjects) PLV was also computed across shorter 2-s data segments (from all blocks) that were time-locked to each partner’s individual movement onsets (see below for details on the algorithm used to identify movement onset time). The resulting PLV was then averaged across time points (Supplementary Figure 1). To avoid edge artefacts, data segmentation was performed on the already band-passed filtered and Hilbert-transformed signals.

Further, to evaluate whether submovements are phase-locked to movement onset, we quantified the within-subject (or inter-movement) PLV on the same data segments as described above but computing the mean resultant vector over the same-participant instantaneous phases (instead of the between-participants phase differences; Supplementary Figure 1).

#### Time-domain analysis

To estimate onset/offset of each individual movement, we low pass filtered (3 Hz, two-pass Butterworth, third order) the position data before computing the velocity. Movement onset was defined as the first data sample of 150 consecutive samples (i.e., 0.5 s) where the velocity was positive (or negative, depending on movement direction); movement offset was defined as the first sample after at least 300 samples (1 s; exact time could be slightly changed according to individual movement duration) from movement onset where the velocity passed through zero. This algorithm was applied iteratively by sliding along the entire velocity time series. Data segmentation was visually checked for each time series and any error was manually corrected (<5%).

The cross-correlation analysis was performed on the raw, unfiltered, data. For each couple and condition, we first took velocity segments corresponding approximately to individual movements, i.e., time-locked to the participant-specific movement onsets and with length equal to mean movement duration (velocity segments belonging to movements with duration > or < than mean duration ± 2.5 SD were discarded from the analysis). Velocity sign was adjusted to be positive in all segments. To remove the movement-locked components, we subtracted from each segment the average velocity profile across all the retained segments (subject-wise). After these common preprocessing steps, we computed the cross-correlation (normalized so that the autocorrelations at zero lag are identically 1) by aligning the two partners’ velocity segments in two different ways: 1) movement-aligned, i.e., keeping the data aligned to each participant’s movement onset as just described, and 2) time-aligned, i.e., using the movement onsets of only one of the two participants (subject ‘A’ for convention) as reference temporal markers to re-align her/his partner’s data (subject ‘B’; Figure 4A,B). Therefore, only the second type of alignment (time-aligned) preserved the real time relationship between the two partners’ velocities. For the ‘Virtual partner’ task, the analysis pipeline was the same, but the cross-correlation was computed between all participants (‘A’ and ‘B’) and the dot velocity, using the latter as the reference signal for re-aligning the participants’ velocities (time-alignment).

For the submovement-locked analysis (Figure 4E, F, G, H), position data were low-pass filtered (4 Hz, two-pass Butterworth, third order) before computing velocity. Preprocessing of velocity data (i.e., segmentation, change of velocity sign, subtraction of the across-segments average) was identical to that already described for the cross-correlation analysis. For one of the two participants (again subject ‘A’ for convention), we identified the submovements as local peaks in the velocity, i.e., data points with values larger than neighbouring values (in each velocity segment). We then segmented the same participant (‘A’) as well as her/his partner (‘B’) velocities based on the identified submovements (from - 0.6 to 0.6 s), providing with ‘submovement-triggered’ averages (Figure 4E). Finally, the submovement-aligned data were also used to estimate the probability of producing a submovement given a submovement produced by one’s partner. Specifically, we counted the number of submovements (local velocity peaks) for each time point (from -0.6 to 0.6 s) in subject ‘B’ velocity segments (aligned to subject ‘A’ submovements) and then divided these numbers by the total amount of analysed velocity segments (Figure 4F). We then averaged the computed probabilities within 36 equally spaced and non-overlapping bins between -0.6 and +0.6 s. The same analysis was performed also for the ‘Virtual partner’ task by aligning the participants’ velocities (both ‘A’ and ‘B’) to the submovements contained in the dot kinematics (Figure 4G, H).

#### Statistical analysis

We statistically evaluated whether movements are executed at a different pace compared to the instructed one by performing one-sample t-tests (against 0.25 Hz) for all conditions (solo, dyad-in-phase/anti-phase) on the F0 peak frequency, i.e., the frequency with maximal power in the velocity power spectrum. For the solo condition, the test was applied on individual parameter estimates (df = 59), whereas for the dyadic conditions the tests were applied on couple-wise across-subjects averages of the parameter estimates (df = 29). The difference in movement rate between the in-phase and anti-phase condition was tested by means of a paired samples t-test (df = 29).

We also computed the Pearson correlation across subjects for all conditions (solo, dyad-in-phase/anti-phase) between, on one side, the F0 peak frequencies and, on the other side, the F1 or (separately) F2 peak frequencies of the individual velocity power spectra. To identify the individual peak frequencies, we used the following criteria. For F0, we just took the frequency with maximal power. For F1 and F2, we sought for local peaks in the power spectrum within frequency ranges comprised between 0.5 and 1.25 Hz and between 1.5 and 4 Hz, respectively. Subjects for which no consistent peak was identified were excluded from the corresponding correlation analysis (F1: 10, 3, 2; F2: 6, 2, 16 excluded subjects out of 60 for solo, dyad-in-phase, dyad-anti-phase, respectively). If more than one peak was identified for a given subject, we included the peak with higher power and frequency closer to the across-subjects average peak frequency (F1: 16, 12, 24; F2: 6, 6, 4 subjects out of 60 for solo, dyad-in-phase, dyad-anti-phase, respectively).

Differences in the (between-subjects) PLV between the in-phase and anti-phase conditions were statistically evaluated by means of conventional paired samples t-tests. The tests were applied separately on the PLV averaged between 0.18 and 0.34 Hz for F0, between 0.55 and 0.99 Hz for F1 and between 1.53 and 3.24 Hz for F2 (ranges defined based on the across-couples range of variation – i.e., min-max – in the corresponding PLV peak frequencies; Figure 3B).

To rule out that the difference between the in-phase and anti-phase condition in the (between-subjects) PLV at F2 depends on corresponding condition differences at F0 and/or F1, we used a data stratification approach. We performed two separate stratifications, one aiming at levelling conditions differences in the mean PLV at F0 (i.e., between 0.18 and 0.34 Hz), and the other in the mean PLV at F1 (i.e., between 0.55 and 0.99 Hz). The distributions across-couples of the PLV at F0/F1 were compiled for the in-phase and anti-phase conditions and binned in 10 equally spaced bins. The number of couples falling in each bin for the in-phase and anti-phase condition was then equated by means of a random subsampling procedure which aims at matching as much as possible the PLV group-level condition averages (as implemented in Fieldtrip, function: ft_stratify, method: ‘histogram’, ‘equalbinavg’). Stratification on F0- and F1-PLV led to the removal of 16 and 17 couples (out of 30), respectively. The condition difference in PLV at F2 (i.e., between 1.53 and 3.24 Hz) after stratification was then statistically evaluated by means of independent samples t-tests. To test whether the modulation of submovement probabilities (obtained for subject ‘B’ as a function of time relative to submovements generated by subject ‘A’) was significantly different from what could be obtained by chance, we created a group-level distribution of surrogate data. More specifically, for each couple we preprocessed data and identified the submovements generated by subject ‘A’ in each data segment following the same analysis steps as already described for the original submovement-locked analysis (see above). We then randomly shuffled data segments for subject ‘B’ (i.e., destroying the temporal association between the two partners’ velocities segments) and used the previously identified time stamps (corresponding to subject ‘A’ submovements) to segment the shuffled data and compute the probability of submovements in subject ‘B’ for each time point (from - 0.6 to 0.6 s; probabilities were then binned over time as already described for the main analysis). We repeated this procedure 1000 times for each couple (and condition) and each time we averaged the result across couples, yielding a group-level surrogate distribution. The p-value of the test is given by the proportion of values of the surrogate distribution that exceeds the original (group-level) submovement probability. The obtained p-values were then corrected for multiple comparisons across time by means of the Bonferroni method. We compared submovement probabilities before vs. after time zero by applying paired samples t-tests on the maximal probabilities computed for all subjects (subjects ‘B’; n = 30) in the respective time intervals (i.e., from -0.6 to 0 s and from 0 to +0.6 s).

Finally, we performed analyses at the couple-level. For each couple, we evaluated the symmetry of both the cross-correlation profile as well as the (subject B) submovement-locked profile by applying paired samples t-tests on the maximal values obtained before vs. after lag/time zero, respectively (Bonferroni correction was applied to correct for the multiple individual tests, n = 30).

## Acknowledgments

This work has been supported by BIAL Foundation – Grant for Scientific Research 2020 to A.T., by Ministero della Salute, Ricerca Finalizzata 2016 - Giovani Ricercatori (GR-2016-02361008) and Ministero della Salute, Ricerca Finalizzata 2018 - Giovani Ricercatori (GR-2018-12366027) to A.D., and by the European Union H2020 - EnTimeMent (FETPROACT-824160) to L.F. The funders had no role in study design, data collection and analysis, decision to publish, or preparation of the manuscript.

## Author contributions

A.T. and A.D. conceived the study; A.T., A.D. and J.L. designed the experiments; A.T., M.E., G.N. and N.P. collected the data; A.T. analysed the data; A.T. wrote the original draft; A.T., A.D., J.L. and L.F. reviewed and edited the manuscript; A.T., A.D. and L.F. contributed to funding and supervision.

## References

1. Repp BH. Sensorimotor synchronization: a review of the tapping literature. Psychonomic bulletin & review. 2005;12(6):969–92.

2. Noy L, Dekel E, Alon U. The mirror game as a paradigm for studying the dynamics of two people improvising motion together. Proceedings of the National Academy of Sciences of the United States of America. 2011;108(52):20947–52.

3. Riley MA, Richardson MJ, Shockley K, Ramenzoni VC. Interpersonal synergies. Frontiers in psychology. 2011;2:38.

4. Oullier O, Kelso JAS. Social coordination from the perspective of Coordination Dynamics. In: Meyers RAe, editor. The Encyclopedia of Complexity and Systems Science. Heidelberg: Springer; 2009.

5. Borroni P, Montagna M, Cerri G, Baldissera F. Cyclic time course of motor excitability modulation during the observation of a cyclic hand movement. Brain research. 2005;1065(1-2):115–24.

6. D’Ausilio A, Novembre G, Fadiga L, Keller PE. What can music tell us about social interaction? Trends in cognitive sciences. 2015;19(3):111–4.

7. Neda Z, Ravasz E, Brechet Y, Vicsek T, Barabasi AL. The sound of many hands clapping. Nature. 2000;403(6772):849–50.

8. Keller PE, Novembre G, Hove MJ. Rhythm in joint action: psychological and neurophysiological mechanisms for real-time interpersonal coordination. Philosophical transactions of the Royal Society of London Series B, Biological sciences. 2014;369(1658):20130394.

9. Repp BH, Su YH. Sensorimotor synchronization: a review of recent research (2006-2012). Psychonomic bulletin & review. 2013;20(3):403–52.

10. Haken H, Kelso JA, Bunz H. A theoretical model of phase transitions in human hand movements. Biol Cybern. 1985;51(5):347–56.

11. van der Steen MC, Keller PE. The ADaptation and Anticipation Model (ADAM) of sensorimotor synchronization. Frontiers in human neuroscience. 2013;7:253.

12. Navas F, Stark L. Sampling or intermittency in hand control system dynamics. Biophys J. 1968;8(2):252–302.

13. Miall RC, Weir DJ, Stein JF. Intermittency in human manual tracking tasks. J Mot Behav. 1993;25(1):53–63.

14. Woodworth RS. Accuracy of voluntary movement. The Psychological Review: Monograph Supplements. 1899;3(3):1–114.

15. Doeringer JA, Hogan N. Intermittency in preplanned elbow movements persists in the absence of visual feedback. Journal of neurophysiology. 1998;80(4):1787–99.

16. Pasalar S, Roitman AV, Ebner TJ. Effects of speeds and force fields on submovements during circular manual tracking in humans. Experimental brain research. 2005;163(2):214–25.

17. Miall RC, Weir DJ, Stein JF. Manual tracking of visual targets by trained monkeys. Behavioural brain research. 1986;20(2):185–201.

18. Roitman AV, Massaquoi SG, Takahashi K, Ebner TJ. Kinematic analysis of manual tracking in monkeys: characterization of movement intermittencies during a circular tracking task. Journal of neurophysiology. 2004;91(2):901–11.

19. Hall TM, de Carvalho F, Jackson A. A common structure underlies low-frequency cortical dynamics in movement, sleep, and sedation. Neuron. 2014;83(5):1185–99.

20. Susilaradeya D, Xu W, Hall TM, Galan F, Alter K, Jackson A. Extrinsic and intrinsic dynamics in movement intermittency. Elife. 2019;8.

21. Craik KJ. Theory of the human operator in control systems; the operator as an engineering system. Br J Psychol Gen Sect. 1947;38(Pt 2):56–61.

22. Neilson PD, Neilson MD, O’Dwyer NJ. Internal models and intermittency: a theoretical account of human tracking behavior. Biol Cybern. 1988;58(2):101–12.

23. Sakaguchi Y, Tanaka M, Inoue Y. Adaptive intermittent control: A computational model explaining motor intermittency observed in human behavior. Neural Netw. 2015;67:92–109.

24. Gawthrop P, Loram I, Lakie M, Gollee H. Intermittent control: a computational theory of human control. Biol Cybern. 2011;104(1-2):31–51.

25. Miall RC. Task-dependent changes in visual feedback control: a frequency analysis of human manual tracking. J Mot Behav. 1996;28(2):125–35.

26. Miall RC, Weir DJ, Stein JF. Visuomotor tracking with delayed visual feedback. Neuroscience. 1985;16(3):511–20.

27. Pereira M, Sobolewski A, Millan JDR. Action Monitoring Cortical Activity Coupled to Submovements. eNeuro. 2017;4(5).

28. Jerbi K, Lachaux JP, N’Diaye K, Pantazis D, Leahy RM, Garnero L, et al. Coherent neural representation of hand speed in humans revealed by MEG imaging. Proceedings of the National Academy of Sciences of the United States of America. 2007;104(18):7676–81.

29. Okano M, Shinya M, Kudo K. Paired Synchronous Rhythmic Finger Tapping without an External Timing Cue Shows Greater Speed Increases Relative to Those for Solo Tapping. Sci Rep. 2017;7:43987.

30. Wolf T, Vesper C, Sebanz N, Keller PE, Knoblich G. Combining Phase Advancement and Period Correction Explains Rushing during Joint Rhythmic Activities. Sci Rep. 2019;9(1):9350.

31. McAuley JH, Marsden CD. Physiological and pathological tremors and rhythmic central motor control. Brain. 2000;123 (Pt 8):1545–67.

32. Loram ID, Gawthrop PJ, Lakie M. The frequency of human, manual adjustments in balancing an inverted pendulum is constrained by intrinsic physiological factors. The Journal of physiology. 2006;577(Pt 1):417–32.

33. Bottaro A, Yasutake Y, Nomura T, Casadio M, Morasso P. Bounded stability of the quiet standing posture: an intermittent control model. Hum Mov Sci. 2008;27(3):473–95.

34. Meyer DE, Abrams RA, Kornblum S, Wright CE, Smith JE. Optimality in human motor performance: ideal control of rapid aimed movements. Psychol Rev. 1988;95(3):340–70.

35. McAuley JH, Farmer SF, Rothwell JC, Marsden CD. Common 3 and 10 Hz oscillations modulate human eye and finger movements while they simultaneously track a visual target. The Journal of physiology. 1999;515 (Pt 3):905–17.

36. Hogan N, Sternad D. Dynamic primitives of motor behavior. Biol Cybern. 2012;106(11-12):727–39.

37. Vince MA. The intermittency of control movements and the psychological refractory period. Br J Psychol Gen Sect. 1948;38(Pt 3):149–57.

38. Craik KJ. Theory of the human operator in control systems; man as an element in a control system. Br J Psychol Gen Sect. 1948;38(Pt 3):142–8.

39. Wolpert DM, Miall RC, Winter JL, Stein JF. Evidence for an error deadzone in compensatory tracking. J Mot Behav. 1992;24(4):299–308.

40. Noy L, Weiser N, Friedman J. Synchrony in Joint Action Is Directed by Each Participant’s Motor Control System. Frontiers in psychology. 2017;8:531.

41. Schmidt RC, Carello C, Turvey MT. Phase transitions and critical fluctuations in the visual coordination of rhythmic movements between people. J Exp Psychol Hum Percept Perform. 1990;16(2):227–47.

42. Reed DW, Liu X, Miall RC. On-line feedback control of human visually guided slow ramp tracking: effects of spatial separation of visual cues. Neuroscience letters. 2003;338(3):209–12.

43. Bourguignon M, Jousmaki V, Dalal SS, Jerbi K, De Tiege X. Coupling between human brain activity and body movements: Insights from non-invasive electromagnetic recordings. NeuroImage. 2019;203:116177.

44. Dumas G, Moreau Q, Tognoli E, Kelso JAS. The Human Dynamic Clamp Reveals the Fronto-Parietal Network Linking Real-Time Social Coordination and Cognition. Cerebral cortex. 2020;30(5):3271–85.

45. Roitman AV, Pasalar S, Ebner TJ. Single trial coupling of Purkinje cell activity to speed and error signals during circular manual tracking. Experimental brain research. 2009;192(2):241–51.

46. Kadmon Harpaz N, Ungarish D, Hatsopoulos NG, Flash T. Movement Decomposition in the Primary Motor Cortex. Cerebral cortex. 2019;29(4):1619–33.

47. Williams SR, Shenasa J, Chapman CE. Time course and magnitude of movement-related gating of tactile detection in humans. I. Importance of stimulus location. Journal of neurophysiology. 1998;79(2):947–63.

48. VanRullen R. Perceptual Cycles. Trends in cognitive sciences. 2016;20(10):723–35.

49. Landau AN, Schreyer HM, van Pelt S, Fries P. Distributed Attention Is Implemented through Theta-Rhythmic Gamma Modulation. Current biology : CB. 2015;25(17):2332–7.

50. Fiebelkorn IC, Kastner S. A Rhythmic Theory of Attention. Trends in cognitive sciences. 2019;23(2):87–101.

51. Jia J, Liu L, Fang F, Luo H. Sequential sampling of visual objects during sustained attention. PLoS Biol. 2017;15(6):e2001903.

52. Tomassini A, Spinelli D, Jacono M, Sandini G, Morrone MC. Rhythmic oscillations of visual contrast sensitivity synchronized with action. The Journal of neuroscience : the official journal of the Society for Neuroscience. 2015;35(18):7019–29.

53. Benedetto A, Spinelli D, Morrone MC. Rhythmic modulation of visual contrast discrimination triggered by action. Proceedings Biological sciences / The Royal Society. 2016;283(1831).

54. Tomassini A, Ambrogioni L, Medendorp WP, Maris E. Theta oscillations locked to intended actions rhythmically modulate perception. Elife. 2017;6.

55. Nakayama R, Motoyoshi I. Attention Periodically Binds Visual Features As Single Events Depending on Neural Oscillations Phase-Locked to Action. The Journal of neuroscience : the official journal of the Society for Neuroscience. 2019;39(21):4153–61.

56. Tomassini A, D’Ausilio A. Passive sensorimotor stimulation triggers long lasting alpha-band fluctuations in visual perception. Journal of neurophysiology. 2018;119(2):380–8.

57. Benedetto A, Binda P, Costagli M, Tosetti M, Morrone MC. Predictive visuo-motor communication through neural oscillations. Current biology : CB. 2021.

58. Benedetto A, Morrone MC, Tomassini A. The Common Rhythm of Action and Perception. Journal of cognitive neuroscience. 2020;32(2):187–200.

59. Schroeder CE, Wilson DA, Radman T, Scharfman H, Lakatos P. Dynamics of Active Sensing and perceptual selection. Current opinion in neurobiology. 2010;20(2):172–6.

60. Benedetto A, Morrone MC. Saccadic Suppression Is Embedded Within Extended Oscillatory Modulation of Sensitivity. The Journal of neuroscience : the official journal of the Society for Neuroscience. 2017;37(13):3661–70.

61. Hogendoorn H. Voluntary Saccadic Eye Movements Ride the Attentional Rhythm. Journal of cognitive neuroscience. 2016;28(10):1625–35.

62. Wutz A, Muschter E, van Koningsbruggen MG, Weisz N, Melcher D. Temporal Integration Windows in Neural Processing and Perception Aligned to Saccadic Eye Movements. Current biology : CB. 2016;26(13):1659–68.

63. Tomassini A, Maris E, Hilt P, Fadiga L, D’Ausilio A. Visual detection is locked to the internal dynamics of cortico-motor control. PLoS Biol. 2020;18(10):e3000898.

64. Crevecoeur F, Kurtzer I. Long-latency reflexes for inter-effector coordination reflect a continuous state feedback controller. Journal of neurophysiology. 2018;120(5):2466–83.

