## Supplementary figures for "Interpersonal synchronization of movement intermittency"

### Supplementary Information

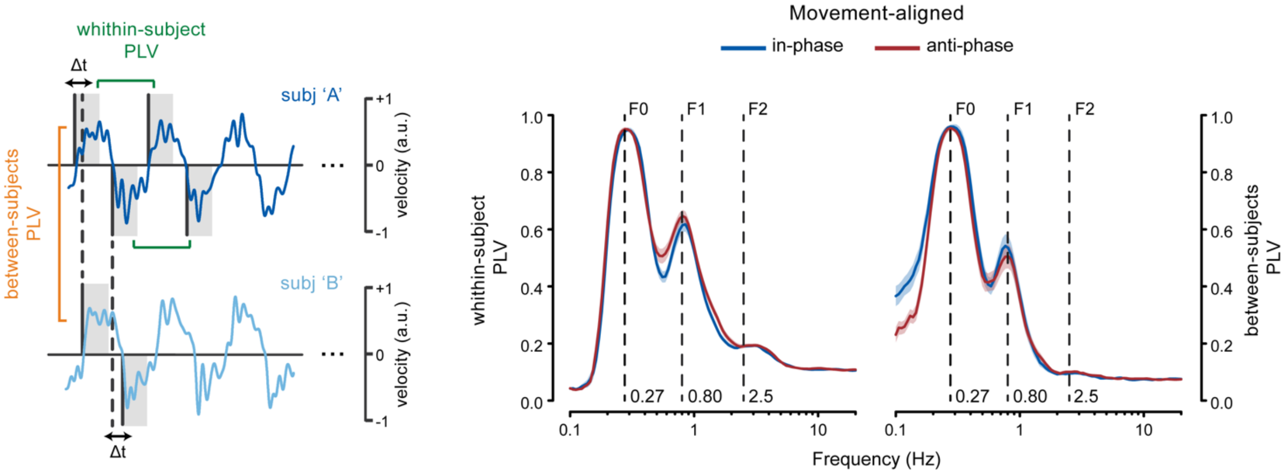


**Figure 1. Movement-locked control analyses: submovement-level interpersonal synchronization is not explained by phase-locking of submovements to movement onset.** The left panel shows example traces of two partners' finger velocity during in-phase synchronization ('Real partner') and a schematic of the data segmentation and analyses. Data are segmented in 2-s segments (corresponding to the instructed movement duration) locked to the onset of single movements performed by each partner. The phase-locking value (PLV) is computed either within-subject (n = 60) – i.e., across the movements performed by the same participant (separately for flexions and extensions and then averaged across movement types) – or between-subjects (n = 30) – i.e., between the two partners' movements. The right panels show the within-subject and between-subjects PLV spectra for both the in-phase and anti-phase conditions (mean ± SEM).

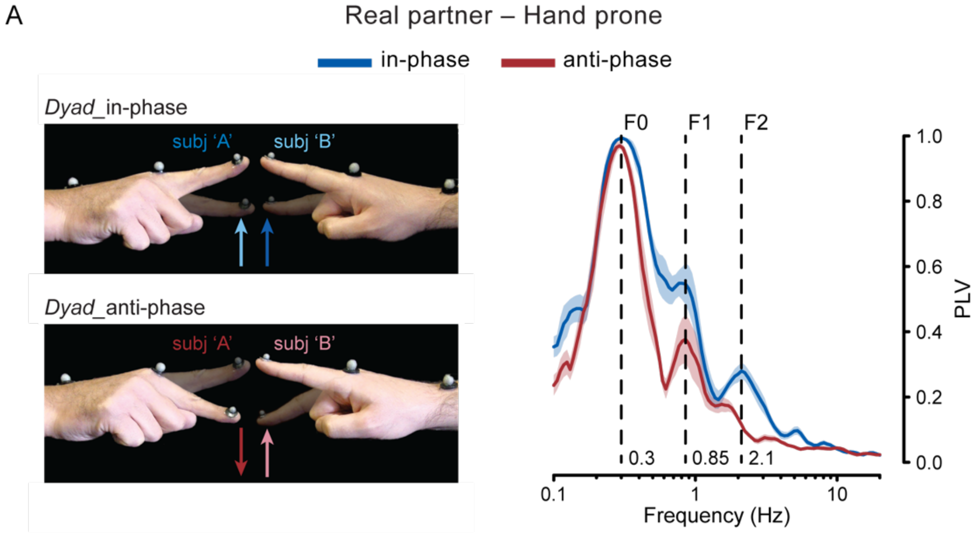


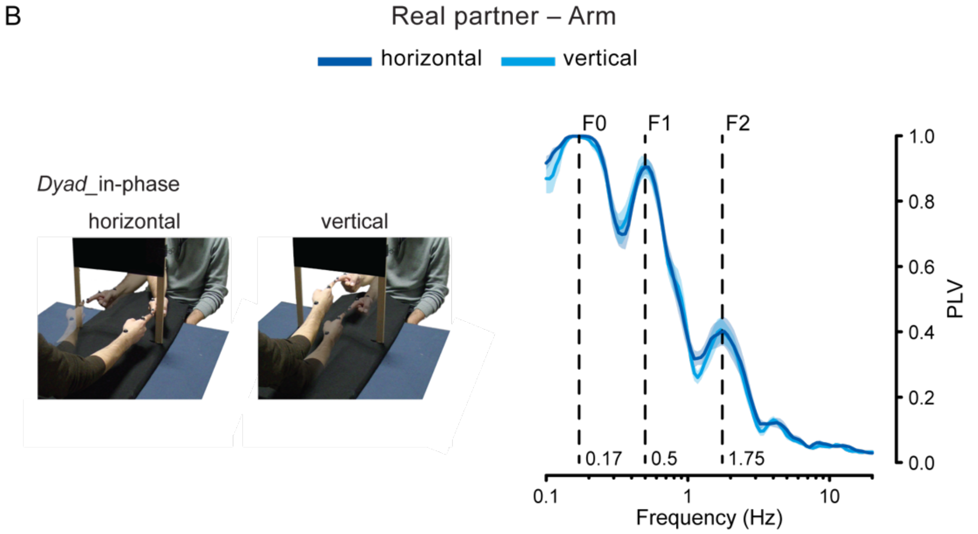


**Figure 2. Secondary tasks: submovement-level interpersonal synchronization does not depend on movement congruency and generalizes to multi-joint actions. (A) left.** Illustration of the 'Real partner – Hand prone' secondary task. Participants keep their right hand in a prone posture and move the index finger along the vertical axis. As opposed to the main task (main Figure 1A, B), the two partners perform simultaneously the same type of movement (either flexion or extension) to attain in-phase coordination (i.e., move towards the same direction), whereas they perform different types of movements (as one performs flexion, the other performs extension and vice versa) to attain anti-phase coordination (i.e., move towards opposite directions). **right.** between-subjects PLV spectra for the in-phase and anti-phase condition (n = 10; mean ± SEM). **(B) left.** Illustration of the 'Real partner – Arm' secondary task. Participants perform whole-arm movements (primarily around the elbow joint) along either the horizontal or vertical inner dimensions of the window (~40x40 cm) delimited by an interposed panel. The task involves tracking of each other's movement to keep the respective fingertips spatially aligned (dyad-in-phase only). Given the larger movement amplitude compared to the main task, the instructed movement rate is reduced to 10 bpm (i.e., ~0.17 Hz). **right.** between-subjects PLV spectra for the horizontal and vertical condition (n = 10; mean ± SEM).
